# Disentangling visual and motor contributions to continuous psychophysics

**DOI:** 10.64898/2026.11.03.763761

**Authors:** Veronica Pisu, Aleksandrs Koselevs, Chloe Lam, Pascal Mamassian, Dominik Straub, Constantin A. Rothkopf, Guido Maiello

## Abstract

Continuous psychophysics measures visual function by asking observers to track a randomly moving target. Because continuous responses provide more information than traditional discrete judgements, a few minutes of tracking can estimate visual sensitivity, making the approach attractive for testing children and patients. Tracking performance is often summarized by cross-correlating the target and cursor velocities: the resulting cross-correlogram peak, lag and width index how strongly, quickly and sharply the observer responds to the target. Tracking, however, requires both seeing the target and moving the cursor towards it, so poor tracking may reflect impaired vision, impaired motor control, or both. Here, we test whether visual and motor contributions to tracking performance can be told apart. Participants tracked a target with a computer mouse while wearing glasses that simulate two levels of visual impairment, using either their dominant or their non-dominant hand. We find that both manipulations alter the cross-correlogram metrics, lowering the peak correlation and delaying and broadening the response, but in different proportions. Visual impairment mainly delays and broadens the response, and does so in proportion to each participant’s loss of visual acuity. The non-dominant hand mainly lowers the peak correlation. Two signatures of motor control explain this difference: with the non-dominant hand, participants pause more often and for longer, and move more noisily between pauses, whereas visual impairment alters neither. Together, response lag, time paused and movement noise reveal whether a change in tracking is visual or motor in almost every participant. Our findings demonstrate that, given a baseline measurement, the same tracking data that measure vision can also reveal whether a change in performance reflects a change in what an individual sees or in how they move.

**Author summary:** We investigate whether it is possible to tell apart the visual and motor contributions to how well people track a moving target. We show that blurring vision with special glasses, and asking people to track with their less skilled hand, both make tracking worse according to standard measures, but not in the same way. Poorer vision mainly slows the response to the target, and the slowing grows with how much the glasses impair each person’s sight. The less skilled hand instead stops moving more often and moves more erratically when it does move, which weakens how closely the cursor follows the target. Poorer vision changes neither of these motor signatures. We then show that these differences reveal, for almost every person, whether their tracking changed because of what they saw or because of how they moved. Our approach could therefore help interpret changes in tracking performance over time when both vision and motor control may change, such as during development, aging and disease.

## Introduction

Psychophysics has revealed much of what we know about perception, yet the dominant paradigms of forced-choice tasks are slow and tedious. Each trial yields roughly one bit of information, and observers must remain engaged over hundreds or thousands of repetitive judgments. Continuous psychophysics overcomes this limitation by replacing discrete trials with a tracking task, in which observers follow a randomly moving target with a cursor or with their eyes [1–6]. This continuous response provides much more information than discrete judgments, such that a few minutes of tracking yield estimates of visual sensitivity that would otherwise require hours of testing [1]. Tracking is also easy, intuitive and engaging, which makes it well suited for testing children, non-human primates and clinical populations [2,5,7–10].

Tracking data are typically analyzed by computing the cross-correlogram between the velocities of the target and the response [1]. When the target follows a random walk, the cross-correlogram describes how the response unfolds over time following a target displacement. The height of the cross-correlogram peak, the lag at which the peak occurs, and the cross-correlogram width all vary lawfully with target visibility. As a target becomes harder to see, the cross-correlogram becomes lower, later, and broader [1,5,11]: the peak decreases while the lag and width increase. These metrics are easy to compute, require no assumptions about the observer, and are closely related to traditional measures of visual sensitivity [1,5].

Tracking, however, is not a purely perceptual task. To follow a target, an observer must estimate its position from noisy sensory signals, plan a movement towards the estimated position, and execute that movement with a noisy and sluggish effector, all while trading off accuracy against effort [12,13]. Any of these stages could limit tracking performance, and the stages interact: gaze tracks a target more accurately when the observer’s own hand moves it, even when vision is blurred [14]. Indeed, Bonnen et al. [1] found that estimates of perceptual uncertainty obtained by fitting a Kalman filter [15] to tracking data were highly correlated with, but systematically larger than, those obtained from an equivalent forced-choice task. Straub and Rothkopf [13] attributed this discrepancy to the motor component of the tracking task and modeled the observer as an optimal controller with separate parameters for perceptual uncertainty, action variability and control costs. Ideal observer analyses have similarly been used to separate early from late sources of noise in tracking [16]. When only the stimulus is manipulated, as in most laboratory studies, the motor contribution can be assumed to remain constant [5]. This assumption may no longer hold if continuous psychophysics is used to track individuals longitudinally. This is the case in many of the settings in which continuous psychophysics appears most promising, such as development, aging, and clinical populations, where both visual and motor function may change over time. If a patient’s tracking deteriorates between assessments, does the patient see worse or move worse?

Here, we test whether visual and motor contributions to tracking performance can be disentangled. We cross three levels of simulated visual impairment with a change of effector, by having participants track a target with either their dominant or their non-dominant hand. This design provides ground truth regarding which system has been altered. We first verify that our visual manipulation impairs visual acuity and contrast sensitivity. We then show that both manipulations make the cross-correlogram lower, later and broader, but in different proportions: simulated visual impairment mainly delays and broadens the response, in proportion to each participant’s loss of acuity, whereas tracking with the non-dominant hand mainly lowers the peak correlation. Finally, we identify two signatures of motor control that account for this difference. With the non-dominant hand, participants pause more and move more noisily between pauses, whereas visual impairment leaves both signatures unchanged. Combined with the lag of the response, these signatures identify the source of a change in tracking performance within individual participants.

## Results

### Simulation glasses impair visual acuity and contrast sensitivity

Fig 1A depicts our experimental design (see Materials and methods for details). Participants used a computer mouse to track a concentric Gabor target that moved following a two-dimensional random walk. Visual impairment was simulated with Cambridge Simulation Glasses [17,18], cardboard glasses with diffusing lenses. Participants wore either no glasses, one pair (low impairment), or two pairs worn one inside the other (mid impairment). Two pairs of glasses impaired both visual acuity (Fig 1B) and contrast sensitivity (Fig 1C) in every participant. One pair reduced contrast sensitivity in every participant, and acuity in all but one, whose acuity was unchanged. Compared to unimpaired viewing, both acuity and contrast sensitivity decreased progressively with the number of glasses worn (Fig. 1 B, C). All three levels of simulated visual impairment differed from each other on both measures (S1 Table).

**Fig 1.**
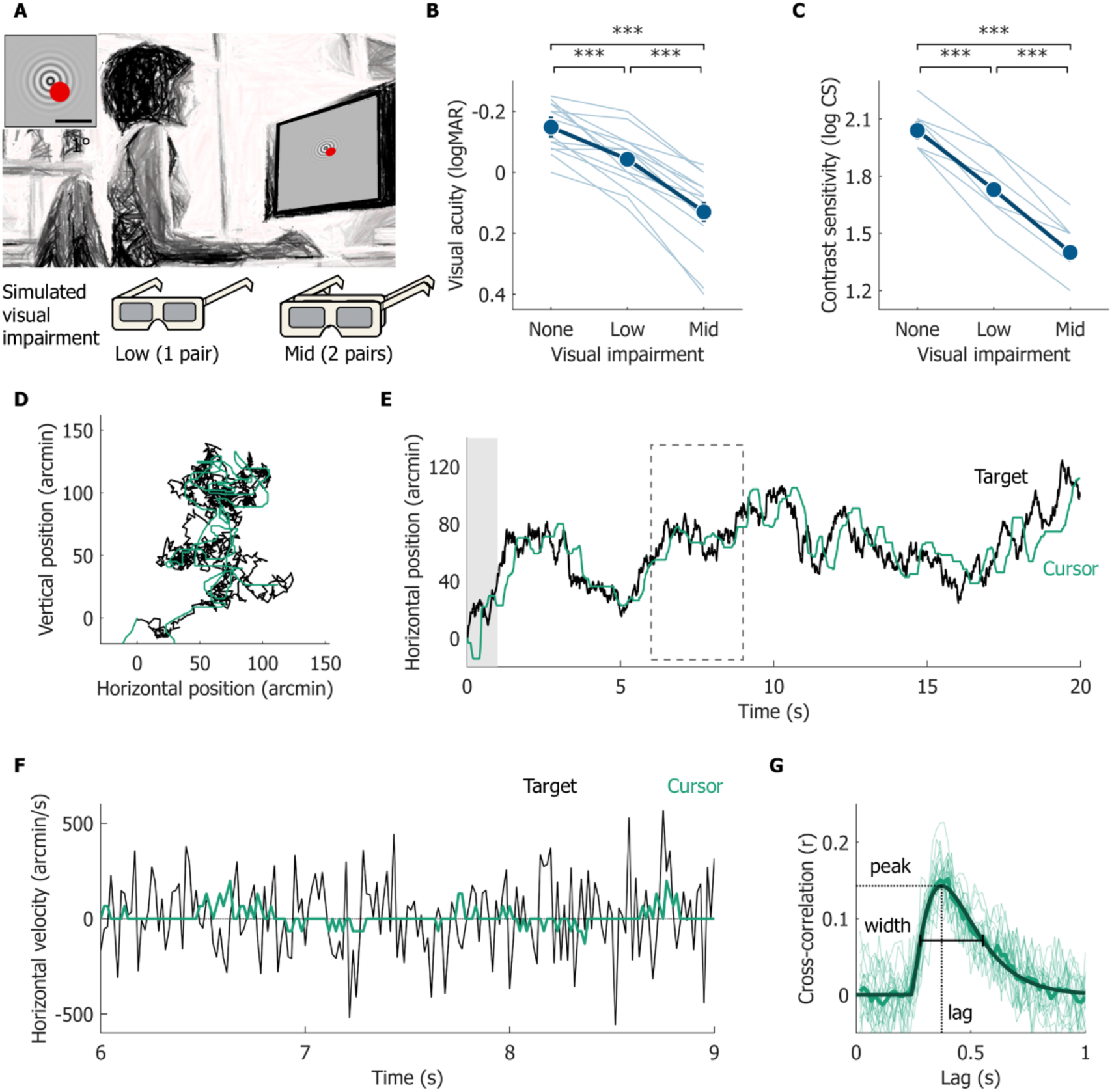
Task, simulated visual impairment and cross-correlogram analysis. (**A**) Participants used a computer mouse with either their dominant or non-dominant hand to track a concentric Gabor target with a red cursor (inset: stimulus to scale; scale bar, 1°). They wore either no glasses, one pair of simulation glasses (low impairment), or two pairs worn one inside the other (mid impairment). (**B**) Visual acuity and (**C**) contrast sensitivity at each level of visual impairment. The acuity axis is inverted, so that worse vision is lower in both panels. Thin lines are individual participants, markers are means, and error bars represent within-participant 95% confidence intervals, which in places are smaller than the markers. Brackets indicate pairwise comparisons (Holm-corrected paired-samples t-tests). (**D**) Target (black) and cursor (green) trajectories in a representative trial (participant 5, no visual impairment, dominant hand, trial 5), whose median cross-correlogram was closest to the group mean in this condition. (**E**) Horizontal position of the target and cursor over time for the same trial shown in D. The first second (gray box) was excluded from analysis, and the dashed box marks the interval shown in F. (**F**) Horizontal velocity of the target and cursor between 6 and 9 s for the same trial shown in D and E. (**G**) Cross-correlograms between cursor and target velocities for the horizontal and vertical components of all 10 trials from the same participant and condition shown in D–F (thin lines), their median (thick line), and the best-fitting delayed gamma function (dark line). Dotted lines mark the peak and lag of the response; the horizontal bar marks its width (full width at half maximum). *** p<.001. N = 15.

### Visual and motor impairment degrade the cross-correlogram in different proportions

Fig 1D shows the target and cursor trajectories from a representative trial, and Fig 1E shows the horizontal component of the same trial over time. The cursor follows the target with a delay of a few hundred milliseconds. Note that the cursor trajectory has a staircase appearance: the cursor often remains still for a fraction of a second while the target keeps moving. The velocities of the target and cursor (Fig 1F) are the signals that we cross-correlated, separately for their horizontal and vertical components, as in Bonnen et al. [1]. Fig. 1 G shows the resulting cross-correlograms for the horizontal and vertical components of all 10 trials from a representative participant in one condition (thin lines), and their median (thick line). The cross-correlogram analysis is described in Materials and methods. For each participant and condition, the median cross-correlogram was fitted with a delayed gamma function (Fig 1G, dark line). The delayed gamma described the cross-correlograms well (median R² = 0.95, range 0.86–0.98). From each fitted function we extracted peak, lag, and width, i.e. full width at half maximum (Fig 1G).

Fig 2A shows the cross-correlograms of all participants in all conditions. Compared with no visual impairment and tracking with the dominant hand, respectively, both visual impairment and tracking with the non-dominant hand made the cross-correlogram lower, later and broader. The effects of the two manipulations were additive, i.e. none of the three metrics showed an interaction between visual impairment and the tracking hand (S2 Table).

**Fig 2.**
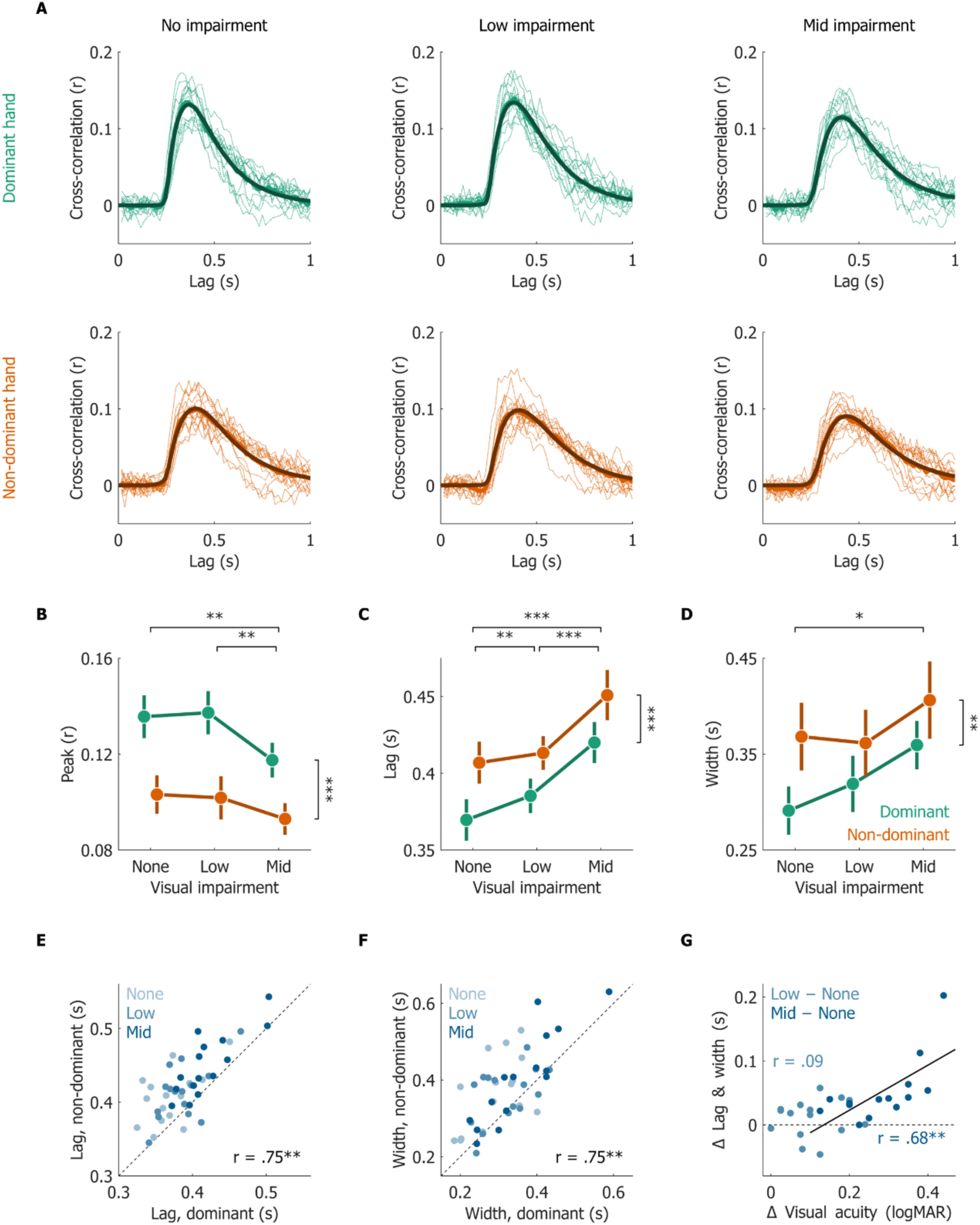
Visual and motor impairment degrade the cross-correlogram in different proportions. (**A**) Cross-correlograms for each condition. Top row: dominant hand (green); bottom row: non-dominant hand (orange); columns: no, low and mid simulated visual impairment. Thin lines are the median cross-correlograms of individual participants, thick lines are their means, and dark lines are the means of the individual best-fitting delayed gamma functions. (**B–D**) Peak (B), lag (C), and width (D) of the fitted cross-correlograms as a function of visual impairment, for the dominant (green) and non-dominant (orange) hand. Markers are means across participants, and error bars represent within-participant 95% confidence intervals. Brackets above the plots indicate pairwise comparisons between levels of visual impairment (collapsed across hands; Holm-corrected paired-samples t-tests); brackets on the right indicate the main effect of hand (repeated-measures ANOVA). (**E, F**) Lag (**E**) and width (**F**) measured with the non-dominant hand against those measured with the dominant hand. Each point is one participant at one level of visual impairment (light to dark blue: none, low, mid), and the dashed line is the identity line. Pearson r between hands is computed on participant means across levels of visual impairment. (**G**) Change in lag and width (mean of the two, averaged across hands) against loss of visual acuity, for low minus no impairment (light) and mid minus no impairment (dark). The line is the least-squares fit to mid minus no impairment, and r values are Pearson correlations for low (light) and mid (dark) minus no impairment. * p<.05, ** p<.01, *** p<.001. N = 15.

The two manipulations, however, differed in their profile (Fig 2B–D). Visual impairment primarily affected the timing of the response, i.e., its lag and width. Mid impairment delayed and broadened the response. Lag was also the only metric to detect the low level of impairment (low vs. no impairment; S2 Table). Tracking with the non-dominant hand, instead, primarily affected the peak correlation.

### Lag and width are consistent across hands and track each participant’s loss of acuity

If the lag and width of the cross-correlogram primarily reflect the visual components of the task, individual differences in these measures should be consistent across hands. Fig 2E and 2F plot, for each participant and level of visual impairment, the lag and width measured with the non-dominant hand against those measured with the dominant hand. Participants who responded later or more sluggishly with one hand also did so with the other, so that the lag and the width of the two hands were strongly correlated across participants. In contrast, the peak correlation was not reliably correlated between the two hands (S3 Table). Lag and width thus capture participant-specific differences that are consistent across effectors, whereas the peak is more specific to the hand used.

We next asked whether the change in response timing (i.e., cross-correlogram lag and width) produced by the glasses reflects how much the glasses impaired each participant’s vision. Lag and width both increased with visual impairment and were consistent across hands. We therefore averaged lag and width to obtain a combined measure, which we then averaged across hands. We chose this combination after observing that the peak did not follow acuity, so this analysis is exploratory. Fig 2G plots each participant’s change in the combined lag-width measure relative to no impairment against their loss of visual acuity. With two pairs of glasses, the response slowed more than with one pair, and greater losses in acuity were associated with greater slowing of the response. This relation held with rank correlation, and after excluding the most influential participant (S3 Table). In contrast, the effect of the hand on lag and width was unrelated to individual loss of acuity. With one pair of glasses, changes in lag and width did not follow the loss of acuity. Two factors limit this comparison: losses of acuity with one pair spanned a narrow range (0.10 ± 0.07 logMAR, mean ± SD; Fig 1B), and the change in lag and width was less reliable across hands than with two pairs (S3 Table).

### The non-dominant hand pauses more often and for longer

Why does tracking with the non-dominant hand lower the peak correlation so much more than visual impairment does? The peak of the cross-correlogram is a correlation coefficient, which decreases whenever the cursor moves in ways unrelated to the target, or fails to move when the target does. Fig 3A shows cursor speed during 3 s of tracking with each hand. With both hands, cursor movements are intermittent: periods of movement alternate with periods in which the cursor remains still, consistent with the intermittent nature of human manual tracking and mouse movements [19–23]. We defined a pause as any period of at least 100 ms in which the cursor did not move along either dimension (i.e., its speed was zero; see Materials and Methods).

**Fig 3.**
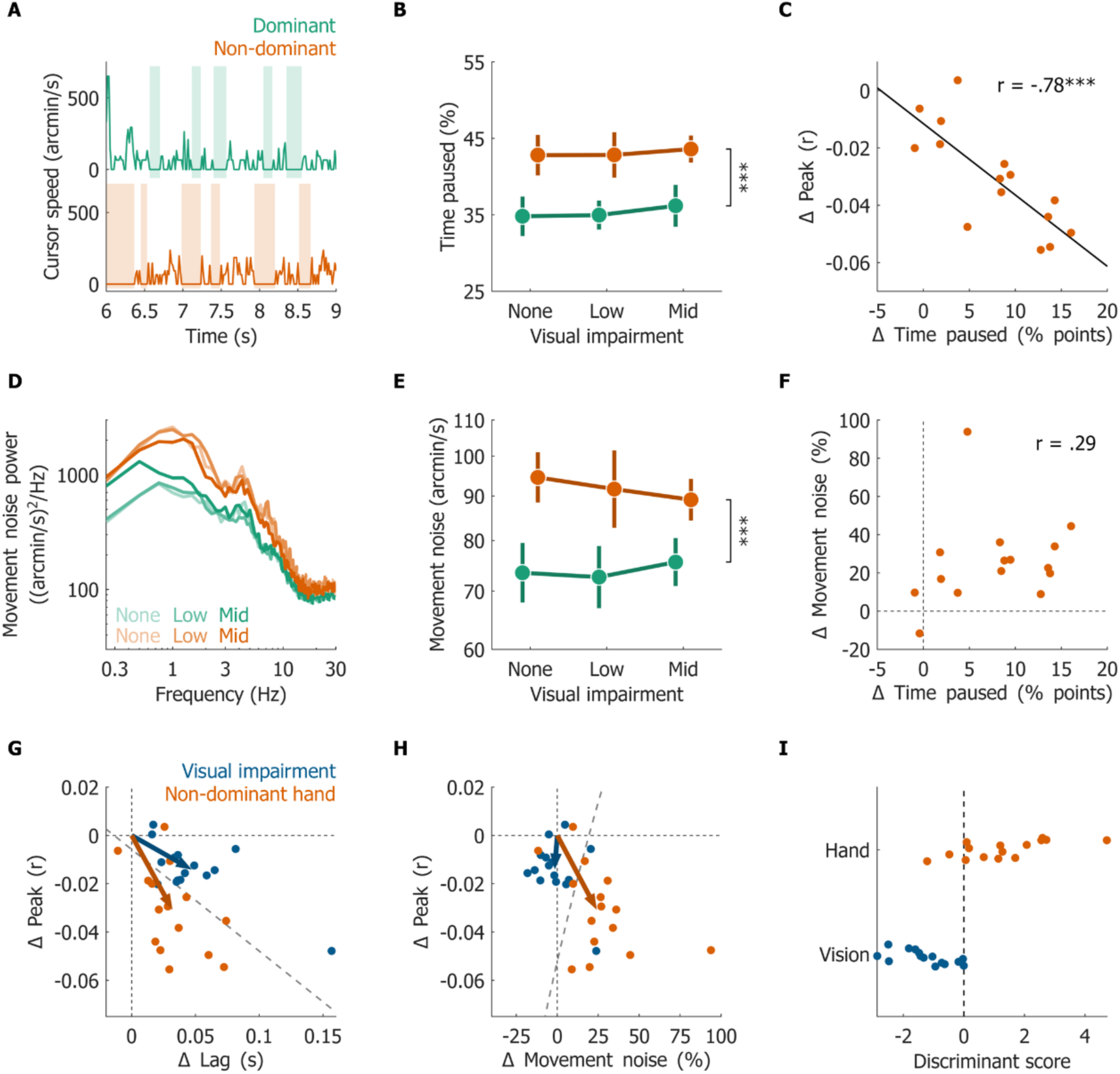
Pausing and movement noise distinguish the non-dominant hand from visual impairment. (**A**) Cursor speed during 3 s of tracking with the dominant (green, top) and non-dominant (orange, bottom) hand, without simulated visual impairment, for the participant with the median effect of hand on time paused (participant 12, trial 1). Shading marks pauses, i.e. periods of at least 100 ms in which the cursor did not move. (**B**) Percentage of time paused as a function of visual impairment, for the dominant (green) and non-dominant (orange) hand. (**C**) Effect of hand (non-dominant minus dominant, averaged across levels of visual impairment) on time paused against its effect on the peak correlation. Each point is one participant, and the line is the least-squares linear regression fit. (**D**) Power spectra of movement noise, i.e. the cursor velocity remaining after accounting for the target-driven response, on frames outside pauses. Lighter to darker lines represent no to mid visual impairment. (**E**) Movement noise as a function of visual impairment (log axis; markers are geometric means). (**F**) Effect of hand (non-dominant minus dominant) on time paused against its effect on movement noise. In C and F, r is the Pearson correlation across participants. (**G, H**) Changes in lag and peak correlation (**G**) and in movement noise and peak correlation (**H**) produced by simulated visual impairment (mid minus none, averaged across hands; blue) and by hand (non-dominant minus dominant, averaged across levels of visual impairment; orange). Each point is one participant, arrows represent mean effects, and the dashed gray line is the linear discriminant boundary fitted to all 30 effects, shown for illustration. (**I**) Cross-validated discriminant score of each effect, from a classifier based on the change in lag, time paused and movement noise for each participant and trained on the other 14 participants. Positive scores are classified as effects of hand; points are jittered vertically for visibility. In B and E, error bars represent within-participant 95% confidence intervals, and brackets on the right indicate the main effect of hand (repeated measures ANOVA). *** p<.001. N = 15.

Participants paused more when tracking with the non-dominant hand than with the dominant hand (Fig 3B). With the non-dominant hand, pauses were more frequent and lasted longer (S4 Table). Simulated visual impairment affected none of these measures. The effect of the hand on time paused was robust to the definition of a pause, and was present for minimum durations between 50 and 200 ms (S5 Table). Importantly, the extent to which each participant paused more with the non-dominant than dominant hand closely tracked how much their peak correlation dropped (Fig 3C). Participants whose non-dominant hand paused more also showed larger increases in lag and width with that hand (S6 Table).

### The non-dominant hand also moves more noisily between pauses

Pauses capture only the moments in which the cursor does not move. We next asked whether tracking with the non-dominant hand also produces more variable movements while the cursor is moving. To characterize cursor movements unexplained by the target, we estimated each participant’s linear response kernel in each condition, i.e. the average cursor velocity following a target step. Convolving the target velocity with this kernel predicts the component of cursor velocity driven by the target. We refer to the remaining cursor velocity, restricted to frames outside pauses, as movement noise (see Materials and methods).

Fig 3D shows the power spectra of movement noise. The non-dominant hand produced more movement noise than the dominant hand across frequencies, whereas spectra measured with the same hand largely overlapped across levels of visual impairment. The effect of the hand was present in every frequency band we examined, from below 0.5 Hz to above 12 Hz, and visual impairment had no significant effect in any band (S5 Table). Accordingly, movement noise (Fig 3E) was 24% greater with the non-dominant hand but did not change with visual impairment. This null effect of vision does not simply reflect a lack of statistical power. Mid impairment changed movement noise by −1.5%, and the 95% confidence interval rules out an increase of more than 5%. A Bayes factor also favored the absence of an effect (S5 Table).

The extra movement noise of the non-dominant hand could, in principle, be a by-product of its extra pausing: after each pause, the cursor must catch up with a target that has kept moving, and these catch-up movements may be less precise. This was not the case. Within the first 200 ms after a pause, the non-dominant hand was 23% noisier than the dominant hand, similar to the difference later in the movement (24%; S5 Table). Moreover, participants who paused more with the non-dominant hand did not reliably show a larger increase in movement noise (Fig 3F), although our sample cannot exclude a moderate relation (S6 Table). Pausing and movement noise thus provide two signatures of motor control, that distinguish the hand manipulation from visual impairment.

### Visual and motor impairments shift tracking performance in different directions

We then asked whether these differences allow the source of a change in tracking performance to be identified within individual participants. For each participant, we computed the effect of simulated visual impairment (mid minus none, averaged across hands) and the effect of hand (non-dominant minus dominant, averaged across levels of visual impairment). Fig 3G plots these effects in the space defined by the change in lag and the change in peak correlation. Simulated visual impairment produced changes mainly along the lag axis, whereas the non-dominant hand produced changes mainly along the peak axis. If the two manipulations affected tracking in the same way, each participant’s visual effect should have no component orthogonal to the mean effect of hand (i.e., the two effects should point in the same direction). Instead, this orthogonal component differed reliably from zero, as did the component of each hand effect orthogonal to the mean visual effect (S6 Table). Movement noise also separated the two effects (Fig 3H): visual impairment lowered the peak without changing movement noise, whereas the non-dominant hand lowered the peak and increased movement noise.

Finally, we asked whether the type of impairment can be identified in individual participants. We used leave-one-out cross-validation to train a linear discriminant classifier on the effects of 14 participants and test it on the effects of the remaining participant. This was repeated so that each participant served as the test participant once. Based on the change in lag, time paused, and movement noise, the classifier correctly labeled 28 of 30 effects (93%) as visual or motor (Fig 3I), where chance is 50%. None of 5,000 random relabelings of the effects achieved this accuracy (S6 Table). We selected these three features after comparing all combinations of the four measures, so this estimate is likely optimistic. Nevertheless, eight of the ten other combinations also classified the effects above chance, with accuracies ranging from 77% to 87% (23–26 of 30). This includes the two cross-correlogram metrics alone, peak and lag (26 of 30), consistent with the different proportions in which the two manipulations affect them. The two combinations that did not differ from chance (20 and 21 of 30) both paired the peak with time paused, whose hand effects were strongly correlated across participants (Fig 3C; S6 Table). Together, these findings show that visual and motor impairments leave distinct signatures in tracking data, which can distinguish the source of a change in performance within individual participants.

## Discussion

We investigated whether visual and motor contributions to continuous tracking performance can be disentangled. Our data show that simulated visual impairment and tracking with the non-dominant hand both altered the cross-correlogram, making the response lower, later, and broader. However, the two manipulations affected these measures in different proportions. Simulated visual impairment mainly delays and broadens the response, with greater losses in acuity associated with greater slowing. Tracking with the non-dominant hand mainly lowers the peak correlation. The non-dominant hand also leaves two signatures that visual impairment does not: participants pause more and move more noisily between pauses. Combined with the lag of the response, these signatures distinguished simulated visual impairment from the effects of tracking with the non-dominant hand in almost every participant. Therefore, although a change in any single cross-correlogram metric is ambiguous, the same tracking data contain the information needed to tell vision and action apart.

### What the cross-correlogram measures

Our findings suggest a simple account of the cross-correlogram metrics. The peak is a correlation coefficient and therefore decreases whenever the cursor fails to move while the target moves, as during a pause, or moves in ways unrelated to the target, as with movement noise. Simulated visual impairment had little effect on either, and thus lowered the peak only modestly. The lag and width of the cross-correlogram, on the other hand, describe the timing of the response. Lag and width increased with visual impairment, were consistent across the two hands, and followed individual loss of acuity. We therefore speculate that lag and width primarily reflect the time needed to localize a degraded target, a sensory limit common to both effectors, whereas the peak additionally reflects how continuously and precisely the effector executes the response. Note that the non-dominant hand also delayed and broadened the response, and that these changes were larger in participants whose non-dominant hand paused more. Lag and width are thus not pure measures of vision either. Bonnen et al. [1] could interpret changes in the cross-correlogram as changes in visual sensitivity because their experiments manipulated only the stimulus. Our results show that this interpretation relies on the motor contribution remaining constant. When the motor system may also change, the cross-correlogram metrics must be interpreted jointly with measures of motor behavior.

### Intermittency and noise in manual tracking

Human manual tracking has long been known to be intermittent, with corrective movements issued at discrete intervals rather than continuously [19–22]. Changing the effector left a clear signature on this intermittency: the non-dominant hand spent roughly a fifth more time paused than the dominant hand, with longer and slightly more frequent pauses. The non-dominant hand also moved more noisily at all frequencies, and was just as noisy later in the movement as right after a pause, when the cursor has to catch up with the target. These findings are consistent with the view that the dominant arm is specialized for controlling limb dynamics [24], with the non-dominant hand executing corrective movements less precisely. Similarly, Mathew et al. [25] found that right-handed participants tracked a moving target less accurately with their left hand and attributed this difference to motor control rather than to prediction. In our experiment, pausing and movement noise were not detectably related across participants, which suggests that pausing and movement noise may reflect partly distinct aspects of motor control. It remains to be seen whether this distinction holds in larger samples.

Intermittency also has implications for model-based approaches to continuous psychophysics. Both the Kalman filter observer of Bonnen et al. [1] and the optimal controller of Straub and Rothkopf [13] assume that the observer responds continuously to every target displacement. Neither model includes pauses in control, and in both, the error between the participant’s tracking position and the model will therefore be absorbed across all noise parameters. Our model-free signatures thus provide empirical constraints for models of the observer: a motor impairment that increases pausing and movement noise, and a visual impairment that does neither. Models of intermittent control, which have been applied to manual tracking and to mouse movements [22,23,26], offer one way to capture these signatures, and may thus yield more trustworthy estimates of perceptual and motor parameters.

### Tracking individual changes in vision

The change in lag and width produced by two pairs of glasses followed each participant’s loss of acuity, measured independently with a letter chart. This suggests that tracking is sensitive not only to group-level manipulations of visibility, but also to individual differences in how much vision is impaired. With one pair of glasses, the change in lag and width did not follow the loss of acuity. The individual losses of acuity with one pair spanned a narrow range, and the change in lag and width was less consistent across the two hands than with two pairs. Testing a wider range of impairments, or repeating measurements within each participant, would be needed to establish the smallest change in vision that tracking can reliably detect in an individual.

### Limitations

We acknowledge some methodological limitations of our work which may be relevant to future research. First, visual impairments were simulated, and the motor manipulation consisted of a change of effector in healthy participants. Clinical visual and motor deficits may leave different signatures. The simulation glasses also reduced acuity and contrast sensitivity together, so our data cannot tell which aspect of vision drives the changes in lag and width. Second, our approach identifies changes relative to an individual’s baseline: time paused and movement noise varied considerably across participants tracking with the same hand, so a single measurement cannot yet establish whether an individual’s vision or motor control is atypical. Third, several analysis choices were made after inspecting the data, namely the combination of lag and width, the features of the classifier, the bounds of the equivalence test and the frequency bands of the spectral analysis. With 15 participants, the accuracy of individual classification is also estimated imprecisely (95% CI: 78–99%). It remains to be seen whether these choices generalize to independent datasets, for example across manual and oculomotor tracking [6,27]. Finally, cursor position was recorded in discrete pixel steps, which may inflate the number of brief pauses; reassuringly, the effect of the hand on time paused was similar for pause definitions between 50 and 200 ms.

### Applications

Our approach may be applied whenever the same individual is tested repeatedly and either vision or motor control may change. Examples include monitoring the treatment of amblyopia [28] or cataract, following the progression of neurological disease [9], assessing visual and motor function in children [8], and measuring the effects of fatigue or medication. Because the analysis requires only the target and cursor trajectories that every continuous psychophysics experiment already records, it can also be applied retrospectively to existing datasets. The same logic should extend to eye tracking, in which oculomotor and visual contributions to tracking performance are similarly entangled [2,5,7–10]. If this were the case, continuous psychophysics could provide not only a rapid measure of visual function, but also a means of determining whether a change in performance reflects a change in what an individual sees or in how they move.

## Materials and methods

### Ethics statement

All procedures were approved by the Psychology Ethics Committee of the University of Southampton (ERGO II, no. 94166). All participants read a participant information sheet and provided written informed consent prior to participating.

### Participants

Fifteen participants (7 female, 8 male; age 27.4 ± 5.7 years, mean ± SD, range 21–41 years, for the 14 participants who reported their age) took part in the experiment. All participants had normal or corrected-to-normal vision. Handedness was self-reported: 14 participants were right-handed and one was left-handed. The dominant hand was defined as the self-reported preferred hand. Participants were students and staff of the University of Southampton, recruited by opportunity sampling and through the department’s participant pool. Individuals with self-reported visual, cognitive or motor impairments were excluded. Participants received course credits or monetary compensation (£8 per hour).

### Apparatus

Stimuli were presented on the 15.6-inch display of a Dell Precision 7540 laptop (34.9 cm wide, 1920 × 1080 pixels, 60 Hz), viewed from a distance of 57 cm, with head position stabilized by a chin rest. At this distance, one pixel subtended 1.10 arcmin (54.7 pixels per degree). The display was gamma-corrected using photometer measurements, and its luminance ranged from 0.17 to 189 cd/m². Participants controlled the cursor with a standard computer mouse (Viglen PXMUSB01 USB optical mouse). Windows pointer speed was set to 10 (the default) and ‘Enhance pointer precision’ was enabled. The experiment was programmed in MATLAB (R2023b) using the Psychophysics Toolbox [29,30].

### Stimuli

The target was a concentric Gabor, i.e. a radial cosine with a period of 11 arcmin (5.5 cycles per degree) multiplied by a Gaussian envelope with a standard deviation of 22 arcmin, presented at full contrast on a uniform mid-gray background (approximately 93 cd/m²). The cursor was a red disk with a diameter of 33 arcmin. On each frame, the horizontal and vertical positions of the target were each displaced by an independent Gaussian step with a standard deviation of 3 arcmin, so that target velocity was white noise along each dimension.

### Simulated visual impairment

Visual impairment was simulated with Cambridge Simulation Glasses [17,18], cardboard glasses with diffusing lenses that reduce acuity and contrast. Participants wore no glasses (no impairment), one pair (low impairment), or two pairs, one inside the other (mid impairment), over their usual optical correction where applicable. For each level of simulated visual impairment, we measured visual acuity with an ETDRS chart (Precision Vision, Chicago, IL) [31,32] viewed at 2 m, and contrast sensitivity with a Pelli-Robson chart (Precision Vision, Chicago, IL) [33]. The average impairment in visual acuity (0.10 and 0.28 logMAR, Fig. 1 B–C) was close to the manufacturer’s specifications for one and two pairs of glasses (0.08 and 0.29 logMAR, respectively [17,18]).

### Design and procedure

The experiment followed a 3 (simulated visual impairment: none, low, mid) × 2 (hand: dominant, non-dominant) within-participant design, with ten 20-second tracking trials per condition. The order of the three levels of simulated visual impairment was randomized for each participant, with hand order randomized within each level. Participants first completed one practice trial with on-screen instructions. At the start of each trial, the cursor was positioned in the upper-left quadrant of the screen and the target at the center. The trial started when participants moved the cursor onto the target at the center of the screen, and lasted 20 s. Participants were instructed to keep the cursor on the target for the duration of the trial. The experiment was self-paced and participants were encouraged to take breaks. Visual acuity and contrast sensitivity were measured in all visual impairment conditions (mid, then low, then no impairment) at the end of the tracking task. The whole session lasted approximately one hour.

### Cross-correlogram analysis

Target and cursor positions were recorded at 60 Hz and converted from pixels to arcmin. The first second of each trial was discarded, and velocities were computed as frame-to-frame differences in position. For each trial and dimension, we computed the normalized cross-correlation between cursor and target velocities separately for the horizontal and vertical dimension, at lags of 1 to 120 frames (17 ms to 2 s). For each participant and condition, the cross-correlogram was defined as the median across the 20 cross-correlations (10 trials × 2 dimensions).

Each cross-correlogram was fitted with a delayed gamma function, which is zero up to a dead time t₀ and then follows a gamma shape with shape k and scale θ, scaled to a peak height a:

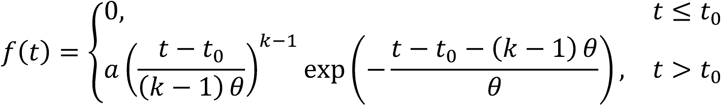

The parameters were bounded (a ∈ [0, 1], t₀ ∈ [0, 0.3] s, k ∈ [1.01, 50], θ ∈ [0.005, 1] s) and fitted by least squares from a grid of starting values, retaining the best fit. Because t₀ and k trade off against each other, we did not analyze the fitted parameters directly. Instead, we extracted the peak, the lag of the peak and the full width at half maximum of each fitted curve, sampled at 0.5-ms resolution. The dead time reached one of its bounds in 4 of 90 fits. Since averaging gamma parameters across participants does not yield a representative curve, the group-level curves in Fig 2A are the means of the individual fitted curves.

### Pauses

A frame was classified as still when the cursor did not move along either dimension. A pause was defined as a run of at least 6 consecutive still frames (100 ms). For each trial, we computed the percentage of time paused, the number of pauses per second and the mean pause duration, and averaged these across the 10 trials of each condition. To assess robustness, we repeated the analysis of time paused with minimum pause durations of 3, 9 and 12 frames (50, 150 and 200 ms).

### Response kernel and movement noise

For each trial and dimension, the response kernel was computed as the cross-covariance between cursor and target velocities at lags of 0 to 120 frames, divided by the variance of the target velocity. For each condition, the kernels were then averaged across the 10 trials and two dimensions to obtain one response kernel for each participant and condition. Because target velocity is white noise, the kernel represents the average cursor response to a unit target step. The target-driven cursor velocity was predicted by convolving the target velocity with the kernel, and the residual was defined as the difference between the measured and predicted cursor velocity. Movement noise was defined as the standard deviation of the residual on frames outside pauses, pooled across the 10 trials and two dimensions and expressed in arcmin/s.

The power spectrum of movement noise was estimated with the Lomb–Scargle periodogram [34,35], which accommodates the gaps left by the pauses. Spectra were estimated at frequencies from 0.25 to 29.75 Hz in steps of 0.25 Hz, averaged across the 10 trials and two dimensions and scaled so that their integral equaled the movement-noise variance. For the spectral analysis, power was averaged within four frequency bands (0.25–0.5, 0.5–3, 3–12 and 12–30 Hz). To test whether extra movement noise reflected catch-up movements, we separately computed movement noise on the frames within 200 ms after the end of a pause and on all later movement frames.

### Statistical analyses

Visual acuity and contrast sensitivity were analyzed with one-way repeated measures ANOVAs. Peak, lag, width, time paused, pause rate, pause duration, log₁₀ movement noise and log₁₀ band power were each analyzed with a 3 × 2 repeated measures ANOVA. All ANOVAs used the Greenhouse–Geisser correction [36], with partial η² as a measure of effect size. Follow-up comparisons between levels of visual impairment (collapsed across hands) and between hands (collapsed across levels of simulated visual impairment) were performed with paired samples t-tests, Holm-corrected for multiple comparisons [37]. Error bars represent Cousineau-Morey within-participant 95% confidence intervals [38,39]. For movement noise, we additionally report the 95% confidence interval of each effect, a default Bayes factor (Cauchy prior on effect size, scale r = .707) [40], a paired comparison between the visual and motor effects, and an equivalence test (two one-sided tests) with bounds of ±8% [41]. The equivalence bounds and the frequency bands were selected after inspecting the data, and these analyses are therefore exploratory.

Agreement between hands was assessed with Pearson correlations across participants, with 95% confidence intervals from the Fisher z-transform. For the relation between the change in lag and width and the loss of visual acuity, we additionally report Spearman rank correlations, the correlation after excluding the participant with the largest Cook’s distance [42], and the split-half reliability of the change across hands (Spearman–Brown corrected).

For each participant, we defined a visual effect (mid minus no impairment, averaged across hands), and a hand effect (non-dominant minus dominant, averaged across levels of visual impairment). Each effect was represented by the changes in two tracking measures (e.g., lag and peak). To test whether the visual and hand effects pointed in the same direction (i.e., whether the two manipulations affected the two measures in the same relative proportions), we projected each participant’s visual effect onto the unit vector orthogonal to the mean effect of hand, and tested the projections against zero with a one-sample t-test. We performed the converse test for the effect of hand, and a leave-one-out version in which the mean direction excluded the tested participant.

To test whether the type of impairment could be identified in individual participants, we used a two-class linear discriminant classifier with pooled covariance. The classifier was trained on the 28 effects of 14 participants and tested on the two effects of the remaining participant. Movement noise entered the classifier as the change in log₁₀ movement noise. Chance performance was assessed by swapping the visual and motor labels within randomly selected participants (5,000 permutations; p computed as the proportion of permutations, including the observed labeling, that performed at least as well). Because accuracy was estimated from only 30 effects, accuracy with shuffled labels varied widely around 50%, with a 95th percentile of 70%. The 95% confidence interval of classification accuracy was computed with the Clopper–Pearson method [43].

## Data and code availability

To facilitate the application of our approach to other datasets, we will provide all raw target and cursor trajectories, the visual acuity and contrast sensitivity data, and MATLAB code reproducing every figure and statistic on a publicly available repository upon journal submission.

## Acknowledgments

Earlier versions of this work were presented as posters at the Vision Sciences Society (VSS) 2025 annual meeting and the European Conference on Visual Perception (ECVP) 2025 and were published as abstracts in the *Journal of Vision* and *Perception*, respectively.

## Declaration of generative AI and AI-assisted technologies in the manuscript preparation process

During the preparation of this work, the authors used Claude (Anthropic) to assist with analysis code, figure drafting, literature searching, and drafting and editing the text. After using this tool, the authors reviewed and edited the content as needed and take full responsibility for the content of the publication.

## Funding

This work was supported by a Fight for Sight Small Grant Award to GM (RESSGA2302; https://www.fightforsight.org.uk) and by a Wessex Medical Trust grant to GM (AF04; https://www.wesmed.org.uk). The funders had no role in study design, data collection and analysis, decision to publish, or preparation of the manuscript.

## Competing interests

The authors have declared that no competing interests exist.

## Author contributions

**Conceptualization:** Veronica Pisu, Guido Maiello, Pascal Mamassian, Dominik Straub, Constantin A. Rothkopf

**Methodology:** Veronica Pisu, Guido Maiello

**Software:** Veronica Pisu, Chloe Lam, Guido Maiello

**Formal analysis:** Veronica Pisu, Guido Maiello

**Data curation:** Veronica Pisu, Guido Maiello

**Visualization:** Veronica Pisu, Guido Maiello

**Investigation:** Veronica Pisu

**Project administration:** Veronica Pisu

**Writing – original draft:** Veronica Pisu, Guido Maiello

**Writing – review & editing:** Veronica Pisu, Aleksandrs Koselevs, Pascal Mamassian, Dominik Straub, Constantin A. Rothkopf, Guido Maiello

**Supervision:** Guido Maiello

**Funding acquisition:** Guido Maiello

**Resources:** Guido Maiello

## Supporting information

**S1 Table.** Visual acuity and contrast sensitivity with the simulation glasses. Estimates for each level of simulated impairment are means ± SD across participants. The effect of visual impairment was tested with a one-way repeated-measures ANOVA (Greenhouse–Geisser corrected; ηp², partial eta squared). Pairwise comparisons are paired-samples t-tests, Holm-corrected. Differences are the first-named level minus the second (e.g., low − none is low impairment minus no impairment): positive differences in logMAR indicate worse acuity, and negative differences in log contrast sensitivity indicate worse contrast sensitivity.

| **Condition or comparison** | **Estimate** | **Statistical test** |
|---|---|---|
| **Visual acuity (logMAR)** |  |  |
| No impairment | −0.148 ± 0.077 |  |
| Low impairment | −0.043 ± 0.091 |  |
| Mid impairment | 0.129 ± 0.129 |  |
| Effect of visual impairment |  | F(1.62, 22.67) = 97.20, p < .001, ηp² = 0.87 |
| Low − none | +0.105 | t(14) = 5.83, p < .001 |
| Mid − low | +0.172 | t(14) = 10.17, p < .001 |
| Mid − none | +0.278 | t(14) = 11.35, p < .001 |
| **Contrast sensitivity (log units)** |  |  |
| No impairment | 2.040 ± 0.111 |  |
| Low impairment | 1.730 ± 0.187 |  |
| Mid impairment | 1.400 ± 0.146 |  |
| Effect of visual impairment |  | F(1.74, 24.33) = 283.04, p < .001, ηp² = 0.95 |
| Low − none | −0.310 | t(14) = −10.02, p < .001 |
| Mid − low | −0.330 | t(14) = −15.20, p < .001 |
| Mid − none | −0.640 | t(14) = −23.48, p < .001 |

**S2 Table.** Cross-correlogram metrics. Omnibus tests are 3 (visual impairment) × 2 (hand) repeated-measures ANOVAs with Greenhouse–Geisser correction (ηp², partial eta squared). Comparisons between levels of visual impairment are paired-samples t-tests collapsed across hands and Holm-corrected. The hand effect is a paired-samples t-test collapsed across levels of visual impairment. Estimates are mean differences, first-named condition minus second. Delayed gamma fits: median R² = 0.947, range 0.857–0.975.

| Effect or comparison | Estimate | Statistical test |
|---|---|---|
| **Peak correlation** |  |  |
| Visual impairment (omnibus) |  | F(1.56, 21.87) = 16.75, p < .001, ηp² = 0.55 |
| Hand (omnibus) |  | F(1, 14) = 44.05, p < .001, ηp² = 0.76 |
| Visual impairment × hand |  | F(1.68, 23.54) = 1.30, p = .286 |
| Low − none | +0.000 | t(14) = 0.06, p = .949 |
| Mid − none | −0.014 | t(14) = −4.65, p = .001 |
| Mid − low | −0.014 | t(14) = −4.31, p = .001 |
| Non-dominant − dominant hand | −0.031 | t(14) = −6.64, p < .001 |
| **Lag (ms)** |  |  |
| Visual impairment (omnibus) |  | F(1.20, 16.82) = 24.14, p < .001, ηp² = 0.63 |
| Hand (omnibus) |  | F(1, 14) = 29.83, p < .001, ηp² = 0.68 |
| Visual impairment × hand |  | F(1.99, 27.92) = 0.59, p = .562 |
| Low − none | +11 | t(14) = 3.12, p = .007 |
| Mid − none | +47 | t(14) = 5.18, p < .001 |
| Mid − low | +36 | t(14) = 4.83, p < .001 |
| Non-dominant − dominant hand | +32 | t(14) = 5.46, p < .001 |
| **Width (ms)** |  |  |
| Visual impairment (omnibus) |  | F(1.62, 22.68) = 5.08, p = .020, ηp² = 0.27 |
| Hand (omnibus) |  | F(1, 14) = 15.21, p = .002, ηp² = 0.52 |
| Visual impairment × hand |  | F(1.79, 25.00) = 1.81, p = .187 |
| Low − none | +11 | t(14) = 0.77, p = .452 |
| Mid − none | +53 | t(14) = 3.04, p = .026 |
| Mid − low | +43 | t(14) = 2.02, p = .126 |
| Non-dominant − dominant hand | +56 | t(14) = 3.90, p = .002 |

To compare the relative effects of hand and visual impairment across the three cross-correlogram metrics, we calculated the ratio of the hand effect to the visual effect (mid minus none) for each metric. A ratio above 1 indicates a larger effect of hand, whereas a ratio below 1 indicates a larger effect of visual impairment. The ratios were 2.21 for peak, 0.68 for lag, and 1.06 for width. These ratios are descriptive and were not tested statistically.

**S3 Table.**
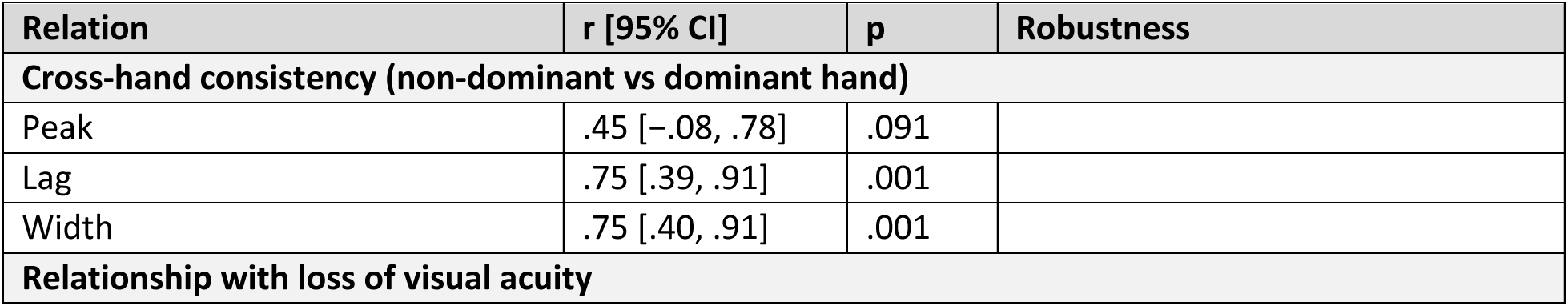
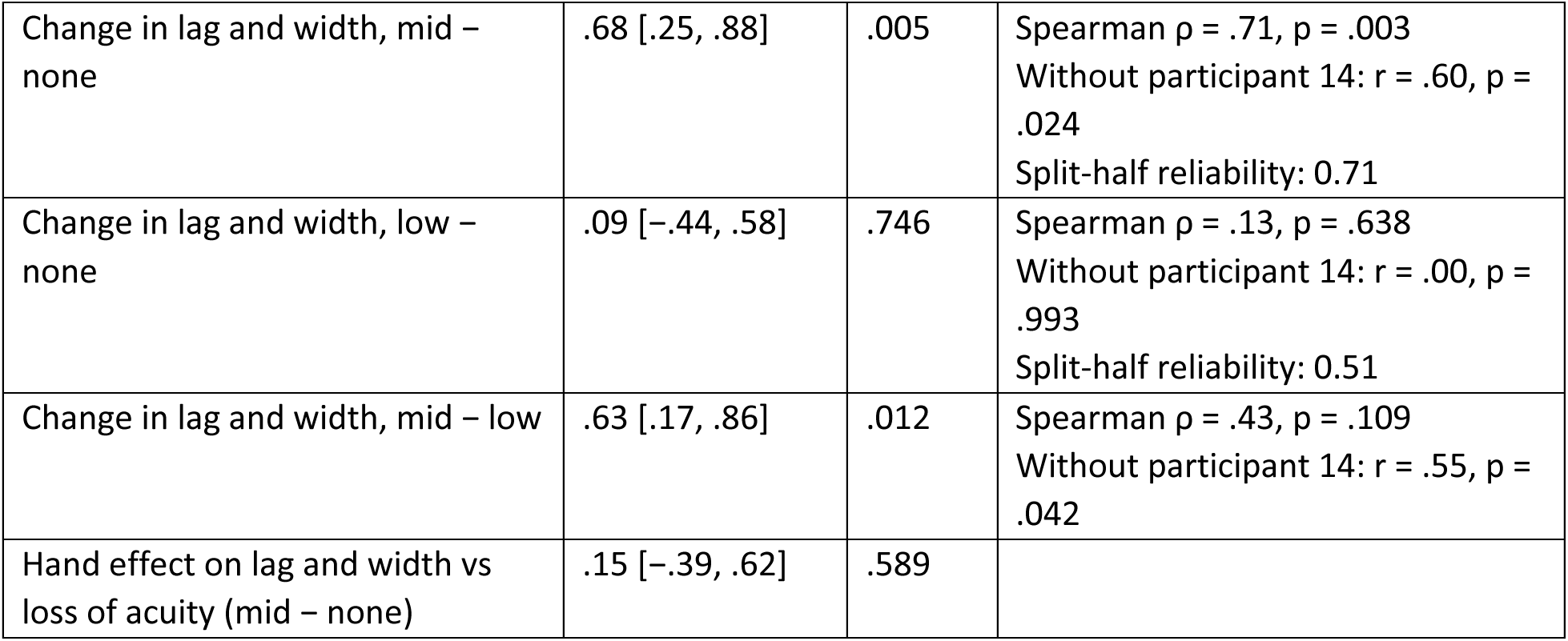
Cross-hand consistency of cross-correlogram metrics and their relationship with visual acuity. Correlations are across participants (Pearson r with 95% CIs from the Fisher z-transform). For cross-hand consistency, measures were averaged across levels of visual impairment. For relationships with acuity loss, changes in lag and width are averaged across hands. Robustness analyses report Spearman rank correlations, exclusion of the most influential participant (i.e., the participant with the largest Cook’s distance), and split-half reliability across hands where applicable. Participant 14 had the largest Cook’s distance (1.69).

**S4 Table.** Effects of visual impairment and hand on pausing and movement noise. ANOVAs as in S2 Table. Movement noise was analyzed as log₁₀ SD; its hand effect is given as the percentage change in SD with 95% CI. The hand effect (non-dominant − dominant) is a paired-samples t-test collapsed across levels of visual impairment. BF₁₀, JZS Bayes factor (Cauchy prior scale 0.707).

| Effect or comparison | Estimate | Statistical test |
|---|---|---|
| **Time paused (%)** |  |  |
| Visual impairment (omnibus) |  | F(1.98, 27.67) = 0.85, p = .437 |
| Hand (omnibus) |  | F(1, 14) = 28.27, p < .001, ηp² = 0.67 |
| Visual impairment × hand |  | F(1.58, 22.12) = 0.05, p = .914 |
| Non-dominant − dominant hand | +7.8 percentage points | t(14) = 5.32, p < .001 |
| **Pauses per second** |  |  |
| Visual impairment (omnibus) |  | F(1.90, 26.53) = 0.40, p = .661 |
| Hand (omnibus) |  | F(1, 14) = 7.73, p = .015, ηp² = 0.36 |
| Visual impairment × hand |  | F(1.95, 27.31) = 1.74, p = .195 |
| Non-dominant − dominant hand | +0.16 | t(14) = 2.78, p = .015 |
| **Pause duration (ms)** |  |  |
| Visual impairment (omnibus) |  | F(1.67, 23.32) = 1.45, p = .253 |
| Hand (omnibus) |  | F(1, 14) = 21.36, p < .001, ηp² = 0.60 |
| Visual impairment × hand |  | F(1.88, 26.39) = 1.28, p = .294 |
| Non-dominant − dominant hand | +26 | t(14) = 4.62, p < .001 |
| **Movement noise** |  |  |
| Visual impairment (omnibus) |  | F(1.84, 25.77) = 0.31, p = .719 |
| Hand (omnibus) |  | F(1, 14) = 24.10, p < .001, ηp² = 0.63 |
| Visual impairment × hand |  | F(1.52, 21.21) = 1.15, p = .322 |
| Non-dominant − dominant hand | +24.2% [13.0, 36.5] | t(14) = 4.91, p < .001, BF₁₀ = 138 |

**S5 Table.** Frequency and robustness analyses of pausing and movement noise. ANOVAs as in S2 Table; band power (Lomb–Scargle) and movement noise were analyzed as log₁₀ values. Percentages are changes in movement-noise SD; hand effects are non-dominant − dominant, collapsed across levels of visual impairment. BF₁₀, JZS Bayes factor (Cauchy prior scale 0.707); BF₁₀ < 1/3 indicates evidence for no effect. The equivalence bounds (±8%) were chosen after inspecting the data.

| Analysis | Estimate | Statistical test |
|---|---|---|
| **Movement-noise power across frequency bands (log<sub>10</sub> power)** |  |  |
| 0.25–0.5 Hz |  | Visual impairment: F(1.48, 20.74) = 1.39, p = .264<br>Hand: F(1, 14) = 49.78, p < .001, ηp² = 0.78<br>Visual impairment × hand: F(1.63, 22.86) = 0.98, p = .376 |
| 0.5–3 Hz |  | Visual impairment: F(1.68, 23.48) = 0.07, p = .902<br>Hand: F(1, 14) = 34.65, p < .001, ηp² = 0.71<br>Visual impairment × hand: F(1.52, 21.28) = 0.83, p = .421 |
| 3–12 Hz |  | Visual impairment: F(1.93, 26.96) = 1.61, p = .219<br>Hand: F(1, 14) = 14.43, p = .002, ηp² = 0.51<br>Visual impairment × hand: F(1.61, 22.50) = 0.56, p = .544 |
| 12–30 Hz |  | Visual impairment: F(1.86, 26.05) = 1.19, p = .317<br>Hand: F(1, 14) = 7.86, p = .014, ηp² = 0.36<br>Visual impairment × hand: F(1.63, 22.88) = 0.26, p = .727 |
| **Movement noise after pauses** |  |  |
| ≤ 200 ms after a pause | Hand: +23.2% | Hand: F(1, 14) = 21.27, p < .001<br>Visual impairment: p = .241 |
| Later in the movement | Hand: +23.7% | Hand: F(1, 14) = 21.32, p < .001<br>Visual impairment: p = .990 |
| **Robustness of the movement-noise effect** |  |  |
| Mid − no impairment | −1.5% [−7.6, +4.9] | t(14) = −0.52, p = .612 |
| Bayes factor, mid − no impairment |  | BF<sub>10</sub> = 0.30 |
| Hand effect vs visual-impairment effect |  | t(14) = 4.79, p < .001 |
| Equivalence test, mid − no impairment within ±8% |  | Two one-sided tests, p = .028 |
| Hand effect, ≤ 200 ms after a pause vs later | +23.2% vs +23.7% | t(14) = −0.16, p = .874 |
| Residual noise over all frames, pauses included (non-dominant − dominant) | +16.3% | Hand: F(1, 14) = 12.26, p = .004<br>Visual impairment: p = .679 |
| **Robustness to pause definition (time paused, dominant vs non-dominant hand)** |  |  |
| Minimum pause duration 50 ms | 46.9% vs 53.0% | Hand: F(1, 14) = 25.36, p < .001<br>Visual impairment: p = .520 |
| Minimum pause duration 100 ms (main analysis) | 35.3% vs 43.1% | Hand: F(1, 14) = 28.27, p < .001<br>Visual impairment: p = .437 |
| Minimum pause duration 150 ms | 26.4% vs 35.0% | Hand: F(1, 14) = 32.40, p < .001<br>Visual impairment: p = .439 |
| Minimum pause duration 200 ms | 19.7% vs 28.2% | Hand: F(1, 14) = 31.92, p < .001<br>Visual impairment: p = .300 |

**S6 Table.** Relationships between motor signatures and separation of visual and hand effects. Correlations are across participants (Pearson r with 95% CIs from the Fisher z-transform) between per-participant hand effects (non-dominant − dominant, averaged across levels of visual impairment). Direction tests are one-sample t-tests against zero of the component of each participant’s effect orthogonal to the mean direction of the other manipulation; visual effects are mid − no impairment, averaged across hands. In the leave-one-out version, the reference direction was computed without the tested participant. Classification used linear discriminant analysis with leave-one-participant-out cross-validation on 30 effects (15 visual, 15 hand); the 95% CI is Clopper–Pearson. Permutation p-values are from 5,000 label swaps within participants, computed as (k + 1)/(5,000 + 1). Chance accuracy is 50%; the 95th percentile of accuracy with shuffled labels was 70%. The primary feature set was selected after comparing all combinations.

| Relationships with time paused (hand effects) |  |  |  |
| Relation |  | r [95% CI] | p |
|---|---|---|---|
| Time paused vs peak |  | −.78 [−.92, −.45] | < .001 |
| Time paused vs lag |  | .62 [.16, .86] | .014 |
| Time paused vs width |  | .67 [.24, .88] | .006 |
| Time paused vs movement noise |  | .29 [−.26, .70] | .288 |
| **Direction tests** |  |  |  |
| **Feature space** | **Tested effects** | **Reference direction** | **Result** |
| Lag and peak | Visual impairment | Mean hand effect | t(14) = 4.58, p < .001<br>Leave-one-out: t(14) = 4.45, p < .001 |
| Lag and peak | Hand | Mean visual effect | t(14) = −5.27, p < .001 |
| Movement noise and peak | Visual impairment | Mean hand effect | t(14) = −3.82, p = .002<br>Leave-one-out: t(14) = −3.82, p = .002 |
| Movement noise and peak | Hand | Mean visual effect | t(14) = 5.35, p < .001 |
| **Classification of effects as visual or motor** |  |  |  |
| **Features** |  | **Correct of 30 (%)** | **Permutation p** |
| Lag + time paused + movement noise (primary) |  | 28 (93%)<br>95% CI [78, 99] | < .001<br>No permutation reached 28 |
| Peak + lag |  | 26 (87%) | .001 |
| Peak + time paused |  | 20 (67%) | .121 |
| Peak + movement noise |  | 24 (80%) | .004 |
| Lag + time paused |  | 23 (77%) | .012 |
| Lag + movement noise |  | 25 (83%) | .005 |
| Time paused + movement noise |  | 25 (83%) | < .001 |
| Peak + lag + time paused |  | 21 (70%) | .057 |
| Peak + lag + movement noise |  | 25 (83%) | .001 |
| Peak + time paused + movement noise |  | 25 (83%) | .003 |
| All four features |  | 26 (87%) | < .001 |

## Notes

### Competing Interest Statement

The authors have declared no competing interest.

